# The impact of transposable elements on tomato diversity

**DOI:** 10.1101/2020.06.04.133835

**Authors:** Marisol Domínguez, Elise Dugas, Médine Benchouaia, Basile Leduque, José Jimenez-Gomez, Vincent Colot, Leandro Quadrana

## Abstract

Tomatoes come in a multitude of shapes and flavors despite a narrow genetic pool. Here, we leveraged whole-genome resequencing data available for 602 cultivated and wild accessions to determine the contribution of transposable elements (TEs) to tomato diversity. We identified 6,906 TE insertions polymorphisms (TIPs), which result from the mobilization of 337 distinct TE families. Most TIPs are low frequency variants and disproportionately located within or adjacent to genes involved in environmental response. In addition, we show that genic TE insertions tend to have strong transcriptional effects and can notably lead to the generation of multiple transcript isoforms. We also uncovered through genome-wide association studies (GWAS) ~180 TIPs associated with extreme variations in major agronomic traits or secondary metabolites. Importantly, these TIPs tend to affect loci that are distinct from those tagged by SNPs. Collectively, our findings suggest a unique and important role for TE mobilization in tomato diversification, with important implications for future breeding.

---

Tomatoes are the highest-value fruit and vegetable crop worldwide. Despite the recurrent genetic bottlenecks that have occurred since its domestication^1,2^, tomato exhibits extensive phenotypic variation and a large part of the diversity seen among the thousands of today’s cultivars likely results from selection of rare alleles with large effects^3^. While genomics-enabled genetics has revolutionized our ability to identify loci underlying domestication and improvement traits in virtually any crop^4–6^, our understanding of the genetic basis of crop diversity is still limited. This situation stems in part from the fact that, with few notable exceptions^7–11^, most genome wide association studies (GWAS) consider only single nucleotide polymorphisms (SNPs) and short indels^12,13^, when structural variants, which include gene presence/absence variants, account for the largest amount of DNA sequence differences between individuals and cultivars^3,10,11,14^. Furthermore, the majority of structural variants result from the mobilization of transposable elements (TEs), which by themselves are potentially an important source of large effect alleles^15^. Indeed, many TEs insert near or within genes^16^ and they have the ability, through the regulatory sequences they harbor, to rewire gene expression networks^16,17^. Although numerous domestication and agronomic traits have been associated with specific TE insertions^15,18–22^, the relative contribution of TEs to the phenotypic diversification of crop species is still poorly documented. Here, we assess through a systematic analysis of 602 re-sequenced genomes the prevalence and impact of TE insertion polymorphisms (TIPs) among wild and cultivated tomatoes.

## RESULTS

### Mobilome composition of tomato

The tomato reference genome (*Solanum lycopersocum* cv. Heinz 1706, release SL2.5) contains 665,122 annotated TE sequences belonging to 818 families^23^. The vast majority of these sequences are ancestral TE copies that have degenerated to varying extent and lost their ability to transpose^24^. To investigate the composition of the tomato mobilome, i.e. the set of TE families with recent mobilization activity, we analyzed short-read whole genome resequencing data available for 602 tomato accessions^2,25,26^. This data set contains wild tomato relatives (Wild; Fig. 1a) and span the Lycopersicon clade, which regroups wild tomatoes (*S. pimpinellifolium*; SP), early domesticated tomatoes (*S. lycopersiucm cerasiform*; SLC) and cultivated tomatoes (*S. lycopersicum lycopersicum* vintage and modern; SLL). To detect additional, i.e. non-reference, TE insertions in each genome sequence, we deployed a refined version of the SPLITREADER pipeline^27^ (Fig. 1b). We restricted our analysis to the 467 TE families with annotated copies longer than 1kb in the reference genome. These families represent the full range of Class I LTR and non-LTR retroelements (i.e. Gypsy, Copia and LINE) and Class II DNA transposons (i.e. MuDR, hAT and CACTA), which move through copy-and-paste and cut-and-paste mechanisms, respectively. After filtering low quality calls (see Methods), 6,906 non-reference TE insertions remained for downstream analysis. Most TE insertions were present in one or a few tomato accessions only (Fig. 1c), suggesting that they occured recently. Nonetheless, cluster analysis based on these 6,906 TIPs recapitulated the phylogenetic relationship between accessions previously determined using SNPs (ref ^2,3^; Fig. 1d).

**Figure 1.**
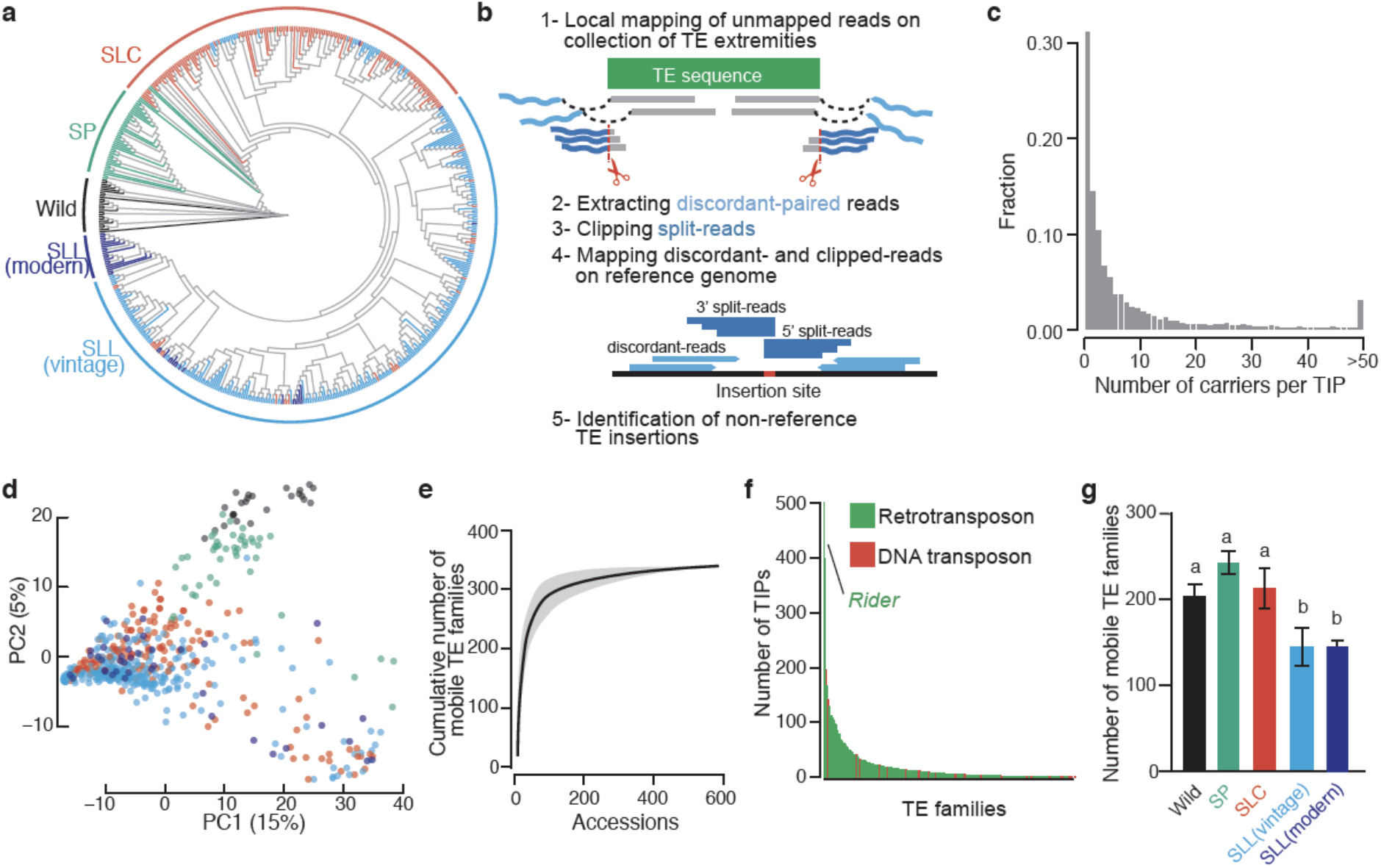
The tomato mobilome. **a.** Phylogeny of the 602 tomato accessions analyzed, including wild tomato relatives (Wild), wild tomatoes (*S. pimpinellifolium*; SP), early domesticated tomatoes (*S. lycopersiucm cerasiform*; SLC) and cultivated tomatoes (*S. lycopersicum lycopersicum* vintage and modern; SLL). **b**. Schematic representation of the SPLITREADER bioinformatics pipeline used to identify TE insertion polymorphisms (TIPs) using split- and discordant-reads. **c.** Distribution of the number of of TIP carriers. **d**. Principal component analysis based on TIPs. Colors represent tomato groups as indicated in a. **e**. Cumulative plot of the number of mobile TE families detected with increasing numbers of accessions. **f**. Number of detected TIPs per TE family. **g**. Number of mobile TE families detected in each tomato group.

TIPs were contributed by 337 TE families, which likely represent the full tomato mobilome, because most TE families with TIPs could be detected using only ~200 of the 602 resequenced genomes (Fig. 1e). The majority (84%) of TIPs resulted from the mobilization of *Ty3/GYPSY* and *Ty1/COPIA* LTR-retrotransposons (Fig. 1f and Supplementary Figure 1a). Notably, the *Ty1/COPIA RIDER* family, which generated insertion mutations with important agronomic implications^21,22,28,29^, contributes the highest number (507) of TIPs overall. Mobilome composition varies substantially among tomato groups and is richest in the genetically diverse SP group (~230 TE families, Fig. 1g). Despite the loss of genetic diversity associated with domestication (ref. ^2,3^ and Supplementary Figure 1b), the mobilome composition of early domesticated SLC is only marginally reduced compared to that of SP (210 vs. 230 TE families, Fig. 1g). This last observation is consistent with the recurrent hybridization between SLC and SP^1^ and the unique ability of TEs to invade new genomes^30^. In contrast, vintage and modern SLL have a more reduced mobile composition (~150 TE families, Figure 1g), consistent with the strong genetic bottleneck caused by the post-Columbian introduction of tomato to Europe.

### TIPs landscape and transcriptional impact

Whereas TE sequences present in the reference genome are enriched in pericentromeric regions^23^, TIPs are distributed more equally along chromosomes (Fig. 2a). Nonetheless, superfamily-specific integration patterns are evident. For instance, TIPs corresponding to *Ty1/COPIA* and many other TE families are found preferentially within or near genes while *Ty3/GYPSY* TIPs cluster in pericentromeric regions (Fig. 2a, b). Importantly, genes harboring TIPs are overrepresented in functions related to response to pathogens or other environmental stresses (Fig. 2c). This overrepresentation is driven by *Ty1/COPIA* LTR-retrotransposons and likely reflects integration preferences rather than relaxed purifying selection or detection biases, which should affect all types of TIPs. Indeed, experimental evidence indicates that *Ty1/COPIA* LTR-retrotransposons integrate preferentially within environmentally responsive genes in *A. thaliana* and rice^31^.

**Figure 2.**
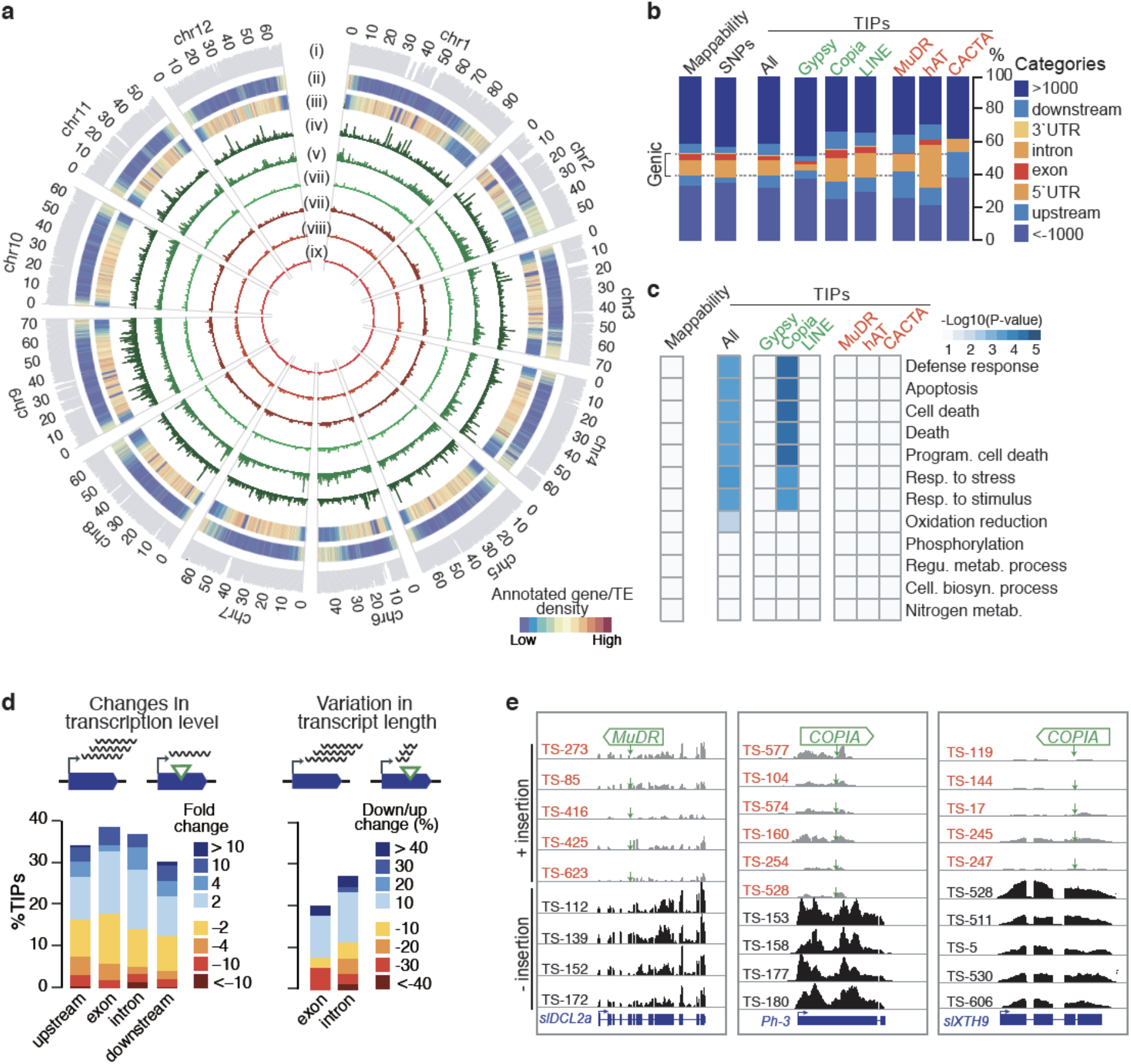
Landscape and transcriptional impact of TIPs. **a.** Chromosomal short-read mappability (i) and distributions of reference genes (ii) and TEs (iii) as well as TIPs by superfamily (iv-ix) across the 12 chromosomes of the tomato genome. (iv) Gypsy, (v) Copia, (vi), LINE, (vii) MuDR, (viii) hAT and (ix) CACTA. **b.** Distribution of TIPs over genic features. UTR, untranslated transcribed region. **c**. GO term analysis of genes with TIPs. **d**. Proportion of TIP-containing genes with changes in transcription level (left panel) or variation in transcript length (right panel) in relation to the presence/absence of the TE insertion. **e**. Genome browser view of RNA-seq coverage for three TIP-containing genes in accessions with (red) or without (balck) the TE insertions. Green arrows indicate the position of TE insertion sites.

In many organisms, including plants and animals, TIPs have been associated with large transcriptomic changes^14,27,32–34^. To assess the impact of TIPs on gene expression in tomato we used RNA-seq data obtained from breaker fruits for 400 accessions^35^. We considered all genes harboring a TIP within 1 kb and compared transcript levels between accessions carrying or lacking the insertion. TIPs associated with two-fold or more changes in gene expression are proportionally more frequent when located in exons and introns (43% and 37%, respectively) than in other gene compartments (Fig. 2d). Furthermore, changes are either positive or negative, consistent with the notion that TE insertions can affect gene expression in multiple ways. To explore further these effects, we compared RNA-seq coverage upstream and downstream of insertion sites (Fig. 2d). This analysis uncovered additional TIPs affecting gene expression and revealed that between 20% and 28% of genic TIPs interfere with transcript elongation when exonic or intronic, respectively. Taken together, these results point therefore to pervasive and complex transcriptional effects of TIPs when these are located within the transcribed part of genes. Consistent with the observed overrepresentation of TIPs within specific gene ontology categories, expression of immune and stress responsive genes was particularly affected by TIPs (Fig. 2e). For instance, we uncovered a rare allele of the gene *slDCL2a* (*Solyc06g048960*) carrying an intronic insertion that is associated with a severe reduction in transcript level. As this gene is involved in resistance against RNA viruses^36^, accessions carrying the rare allele may be more susceptible to viral attacks. Similarly, an exonic *COPIA* insertion within the CC-NB-LRR gene *Ph-3* (*Solyc09g092310*), which confers broad resistance to *Phytophthora infestans*^37,38^, is associated with transcript truncation and might therefore cause increased susceptibility to this pathogen. In addition to such insertions, we also identified TIPs with potential beneficial effects. For instance, an exonic insertion in the gene *slXTH9* (*Solyc12g011030*), which encodes a xyloglucan endotransglucosylase/hydrolase preferentially expressed during fruit ripening^39^, is associated with a near complete loss of expression. Given the key role of *slXTH9* in fruit softening^24^, this natural loss-of-function allele could potentially be harnessed to breed tomato fruits with harder texture and longer shelf life^40^.

### TIPs as an unregistered source of phenotypic variants

To assess more systematically whether TIPs are a potentially important source of phenotypic variation, we first measured the proportion of TIPs in high linkage disequilibrium (LD, *r*^2^>0.4) with SNPs. This proportion was much lower than for SNPs in high LD with other SNPs (Fig. 3a). In agreement with previous findings in Arabidopsis^41^, maize^11^, grapevine^10^ and humans^14^, rare TIPs (MAF<1%) tend to have lower LD with nearby SNPs than more common TIPs (Supplementary Figure 2a,b). In addition, most TIPs in high LD with SNPs arelocated on chromosome 9 (Supplementary Figure 2c), consistent with modern tomatoes harboring on that chromosome a large introgressed segment from wild tomatoes^2^. Based on these observations, we carried out genome-wide association studies specifically using TIPs^9^ (TIP-GWAS, see Methods) for 17 important agronomic traits in tomato, including determinate or indeterminate growth, simple and compound inflorescences, leaf morphology, fruit color, shape and taste (Supplementary Fig 3a). Our TIP-GWAS revealed a total of 22 loci associated with eight traits, including fruit color and leaf morphology (Fig. 3a and Supplementary Figure 3c). These two traits were previously linked to TE insertions^28,42^, thus validating the TIP-GWAS approach. Furthermore, most of these associations could not be identified using SNPs (Supplementary Figure 3b), demonstrating the interest of considering TIPs in addition to SNPs in GWAS. For instance, our TIP-GWAS revealed a strong association between a LTR-retrotransposon *RIDER* insertion within the gene *PSY1*, which encodes a fruit-specific phytoene synthase, and yellow fruit (Fig. 3b,c). Incidentally, SNP-GWAS revealed another variant of *PSY1* associated with yellow fruit (Fig. 3b). Local assembly using Illumina short-reads indicated that this alternative allele, which we named *r^Del^* to distinguish it from the previously identified *r^TE^* allele, contains a ~6kb deletion that bridges the last exon of *PSY1* with the next gene (*Solyc03g031870*) downstream (Fig. 3c). Together, *r^TE^* and *r^Del^* account for 60% of yellow tomato accessions (Fig. 3d,e) and those carrying the *r^TE^* allele display lower expression levels of *PSY1* and yellower fruit than accessions with the *r^Del^* allele (Fig. 3f). Moreover, we detected the *r^TE^* and *r^Del^* alleles in several SLC and SLL vintage accessions but in none of the wild tomatoes (*S. pimpinellifolium*) and wild relatives (Fig. 3g), which suggests that *r^TE^* and *r^Del^* arose after domestication. Also, while the LTR-retrotransposon *RIDER* insertion affected a common haplotype of *PSY1* shared among early domesticated and improved tomatoes, the ~6kb deletion affected a rare haplotype containing numerous SP-derived sequences (Supplementary Figure 4). Together, these results suggest that the first tomato cultivars introduced in Europe during the 16th century, which were reported to be yellow^43^, carried the *r^TE^* allele.

**Figure 3.**
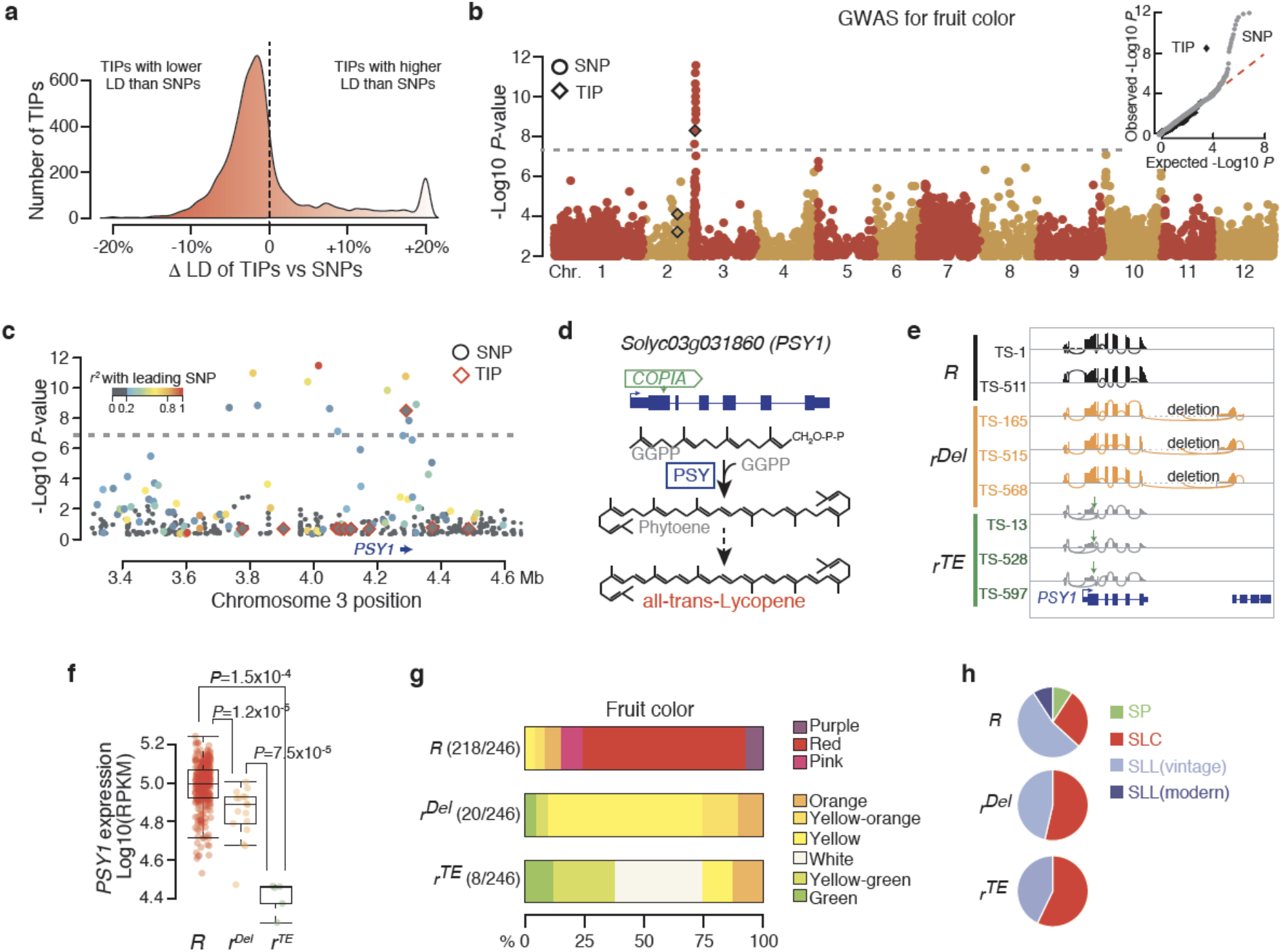
TIPs as an unregistered source of phenotypic variants. **a**. Distribution of the proportion of SNPs that are in lower or higher linkage disequilibrium (LD) with TIPs or other SNPs. **b**. Manhattan plot of SNP- and TIP-based GWAS (circles and diamonds respectively) for fruit color. Inset plot shows the observed and expected distribution of P-values for SNP- and TIP-GWAS (grey circles and black diamonds, respectively). **c**. Manhattan plot of SNP- and TIP-based GWAS (circles and diamonds respectively) around PSY1. Colors indicate the linkage disequilibrium (*r^2^*) with the leading variant. **d**. Structure of the *PSY1* gene with the position of the *RIDER* insertion and simplified representation of lycopene biosynthesis. **e**. Genome browser view of RNA-seq coverage over *PSY1* of accessions carrying the wild-type (*R*) or mutant alleles (*r^del^* and *r^TE^*) of *PSY1*. **f**. Quantification of *PSY1* expression. Statistical significance for differences was obtained using the MWU test. **g**. Fruit color of accessions carrying the distinct alleles of *PSY1.* **h**. Distribution of *PSY* alleles between tomato groups.

To investigate further the specific contribution of TIPs to trait variation in tomato we conducted SNP- and TIP-GWAS on 1,012 metabolic phenotypes measured for more than 397 accessions^25,35^. We identified a total of 846 loci associated with variation for 369 metabolites using SNP-GWAS and 157 loci associated with variation for 60 metabolites using TIP-GWAS (Fig. 4a). There is little overlap between the two sets of loci, thus not only confirming the low LD between SNPs and TIPs but more importantly indicating also that TIPs tend to affect specific sets of genes and traits. As a matter of fact, TIPs unlike SNPs are more often associated with variation in secondary metabolites (Fig. 4c). This enrichment is primarily driven by a higher proportion of TIPs associated with glycoalkaloids and volatiles (Supplementary Figure 5), two classes of compounds that are implicated in defense response and/or interaction with other organisms^44,45^. Also, loci associated with multiple metabolic phenotypes, i.e. association hotspots, are more frequently found with TIPs than SNPs (7% vs. 3%, respectively; *P*<0.01; Fig. 4d). Of note, one such hotspot spans *PSY1* and concerns 15 metabolites, including apocarotenoid volatiles, which are major determinants of tomato flavor (Supplementary Figure 6). In addition, effect sizes are much larger for TIPs than for SNPs (Fig. 4e), consistent with the notion that TE insertions affecting genes tend to create major effect alleles and suggesting that such alleles are better tolerated within secondary metabolism pathways. Finally, almost all TIPs detected by GWAS are present in SLC accessions (Fig. 4f), indicating that their contribution to phenotypic diversification is higher among early domesticated tomatoes.

**Figure 4.**
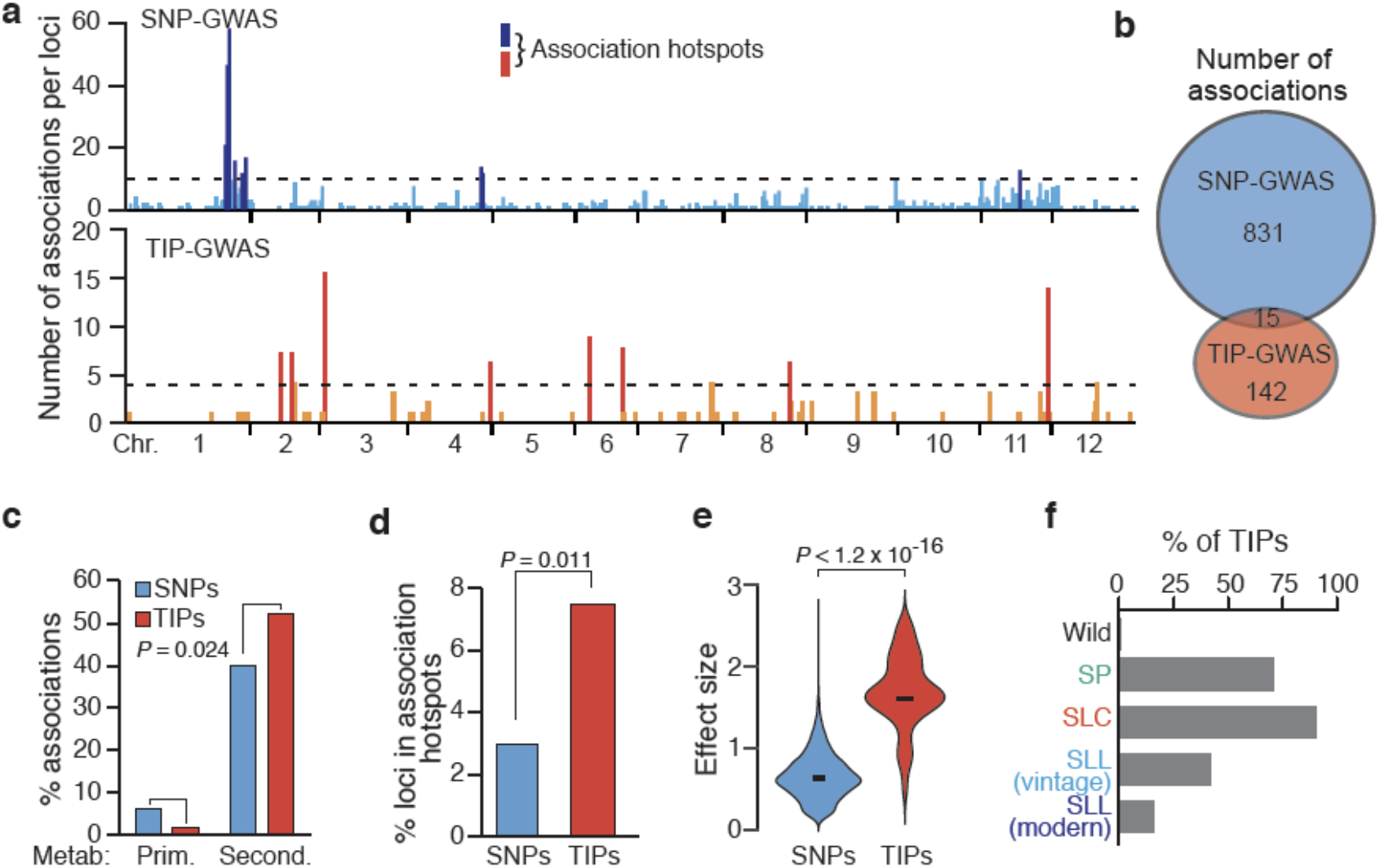
TIPs associations with secondary metabolism. **a.** Distribution of SNP- and TIP-GWAS signals across the tomato genome. **b**. Significant associations detected by SNP- and TIP-GWAS and their overlap. **c.** Percentage of loci statistically associated with primary (Prim.) and secondary (Second.) metabolites. Statistical significance for differences was obtained using the Fisher test. **d**. Percentage of association signals within association hotspots. Statistical significance for differences was obtained using the Fisher test. **e**. Effect size for association signals detected in SNP- and TIP-GWAS. Statistical significance for differences was obtained using the MWU test. **f**. Percentage of TIPs with significant associations present within each of the five tomato groups.

### A key TIP for tomato flavor

Our TIP-GWAS revealed a *COPIA* LTR-retrotransposon insertion that is absent in modern cultivars and associated with high levels of 2-phenylethanol (Figure 5a-c), which gives a pleasant flowery aroma to heirloom tomatoes^46^. This TIP is located in the single intron of gene *Solyc02g079490*, which is preferentially expressed in ripe fruits and encodes a protein with high similarity (63% aa identify) with a cinnamyl alcohol Acyl-CoA transferase^47^ (Supplementary Figure 7a,b). Consistent with a potential role of *Solyc02g079490* in the accumulation of 2-phenylethanol, the introgression line (IL) 2.3^48^, which harbors the lowly expressed *S. pennellii* allele of *Solyc02g079490*^49^, also accumulates more 2-phenylethanol compared to the modern cultivar M82 (ref. ^50^; Supplementary Figure 7c). Thus, *Solyc02g079490* likely encodes a putative 2-phenylethanol Acyl-Coa transferase (PPEAT) involved in the esterification of 2-phenylethanol (Fig 5d), which otherwise accumulates in fruits.

**Figure 5.**
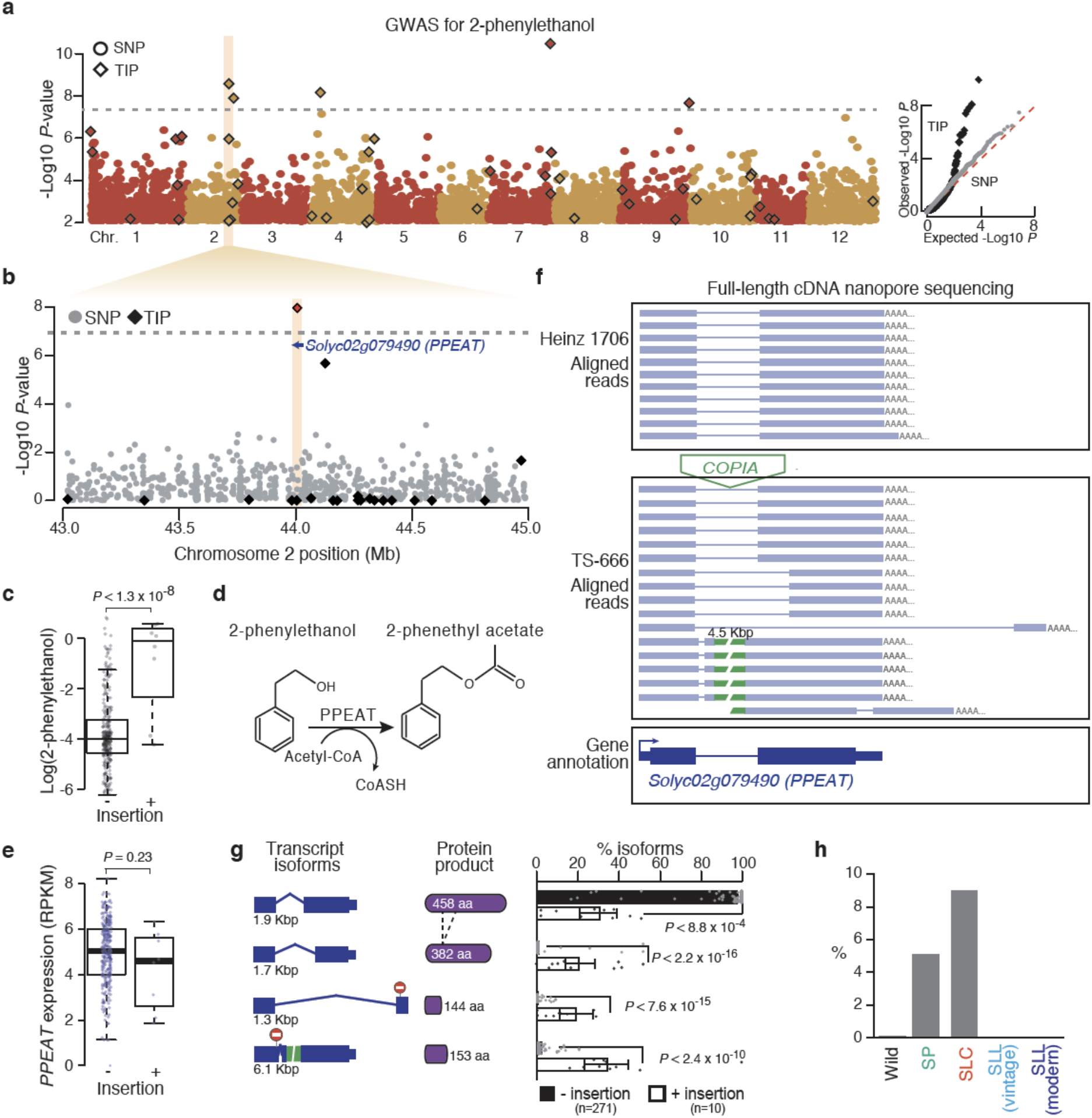
A key TIP for tomato flavor. **a.** Manhattan plot of SNP- and TIP-based GWAS (circles and diamonds respectively) for 2-phenylethanol. qq-plot depicting observed and expected distribution of P-values for SNP- and TIP-GWAS (grey circles and black diamonds, respectively) is shown on the right panel. **b**. Detailed view of the manhattan plot for 2-phenylethanol around *Solyc02g079490*. **c**. 2-phenylethanol levels in accessions carrying or not the intronic *COPIA* insertion. Statistical significance for differences was obtained using the MWU test. **d**. Inferred enzymatic activity of *Solyc02g079490* gene product. **e**. *PPEAT* expression level in accessions carrying or not the intronic *COPIA* insertion. Statistical significance for differences was obtained using t-test **f**. Genome Browser view of full-length cDNA Nanopore reads obtained from two accessions carrying or not the intronic *COPIA* insertion. **g**. *PPEAT* transcript isoforms and protein products. Abundance of transcript isoforms is shown on the right panel. Statistical significance for differences was obtained using the MWU test. **h**. Frequency (%) of the intronic *COPIA*-containing allele in each of the five tomato groups. PPEAT: Putative phenylethanol acyl transferase.

Although the intronic *COPIA* LTR-retrotransposon insertion does not appear to affect the expression levels of *Solyc02g079490* (Fig. 5e), hereafter referred to as *PPEAT*, we noted numerous transcript isoforms in accessions carrying the insertion compared to a single predominant transcript otherwise (Supplementary Figure 7d). To characterize these additional transcripts further we performed full-length cDNA Nanopore sequencing of ripe fruit samples from two accessions, one carrying and one lacking the intronic *COPIA* insertion. We uncovered in this way at least three additional transcript isoforms, all of which result from alternative splicing (Fig. 5f). Moreover, the *COPIA*-containing intron (>5 Kbp), which is one of the largest intron genome-wide based on our Nanopore sequencing (Supplementary Figure 7e), is spliced out in most cases. Nonetheless, this large intron is retained in one isoform, thus leading to an unusually long transcript (Fig. 5f and Supplementary Figure 7e). All of the alternative isoforms incorporate premature stop codons or encode proteins that lack highly conserved catalytic domains (Fig. 5g and Supplementary Figure 7f). Based on these results, we re-analyzed the RNA-seq data obtained for 400 accessions. This new analysis confirmed that truncated isoforms are exclusively associated with the intronic *COPIA* insertion and indicated in addition that these isoforms make up around 60% of all PPEAT transcripts (Fig. 5g). Although a formal demonstration would require the removal of this *COPIA* insertion by genome editing, we can reasonably conclude that it generated a hypomorphic *PPEAT* allele, thus explaining the overaccumulation of 2-phenylethanol. Moreover, because the *COPIA*-containing allele is absent from wild relatives but present respectively at low and intermediate frequency in wild (*S. pimpinellifolium*) and SLC tomatoes (Fig. 5h), we can speculate that it predated domestication and was selected in early domesticated tomatoes but not in modern varieties, which are notorious for their poor flavor^46^.

## Discussion

Cultivated tomato has a complex history of domestication and improvement, characterized by two successive genetic bottlenecks, followed since modern breeding by several introgression events from wild tomatoes and relatives to replenish the limited pool of disease resistance genes^1,10,51^. Despite a relatively narrow genetic diversity, the 25,000 cultivars grown around the world today exhibit an extraordinary phenotypic diversity and the underlying allelic variants are being progressively identified thanks to the advent of high-throughput genome sequencing. In this context, we show that TIPs, which to date have been ignored from population genomic studies in tomato, are an important diversifying force to consider, as has been proposed for other plant species^27,33,34,52–57^. In particular, most TE insertions are low frequency variants, they are typically not tagged by SNPs, representing a previously uncharacterized source of phenotypic variation. Moreover, TIPs tend to affect specific genes, in part as a result of TE insertion preferences, and to associate with larger phenotypic effects. These observations suggest that, in marked contrast to SNPs, most TIPs identified in GWAS as leading variants are likely causal.

Our findings also indicate that TEs have a unique capacity to affect genes involved in environmental response or associated with secondary metabolism. This distinctive property of TEs could be harnessed to breed tomato fruits with superior traits. Moreover, given the limitations associated with the detection of TIPs using short-read sequencing technologies^58^ (i.e. reduced sensitivity and specificity), the analyses presented here certainly underestimates their influence on phenotypes. The development of long-read sequencing technologies should soon provide the means to reveal the full extent of the contribution of TIPs and other structural variants, perhaps mediated by TE sequences, to crop diversity.

The composition of the tomato mobilome, as defined here, appears to be substantially reduced following the post-Columbian introduction of tomato to Europe. This observation may reflect the strong genetic bottleneck this introduction created as well as the increased levels of inbreeding that ensued, which favored the accumulation of deleterious mutations and should thus have compromised the long-term survival of accessions with high mobilome activity^15^. Whether introgression from wild germplasms used in modern breeding can alter this picture by enabling new TE mobilization remains to be determined.

## METHODS

### Detection of TIPs

Illumina re-sequencing data from 602 tomato accessions was downloaded from EBI-ENA and aligned to the tomato genome reference (version SL2.5) using Bowtie2 (arguments –mp 13 – rdg 8,5 –rfg 8,5 –very-sensitive) and PCR-duplicates were removed using Picard. The detection of TIPs was performed using an improved version of SPLITREADER (Baduel et al, Methods in Mol. Biol., *in press*). Compared to the previous version of our SPLITREADER pipeline^27^, we now use the information of both split-reads and discordant reads to call non-reference insertions. In addition, we genotyped the absence of non-reference TE insertions by analyzing local coverage around the insertion sites. SPLITREADER has four steps: (i) extraction of reads mapping discordantly or not at all to the reference tomato genome; (ii) mapping to a collection of reference TE sequences and selection of the reads aligning partially or discordantly; (iii) re-mapping selected reads to the reference genome sequence; (iv) identification of cluster of split-reads and/or discordant-reads indicating the presence of a non-reference TE insertions. Specifically, for each tomato accession we retrieved reads that did not map to the reference genome sequence (containing the SAM flag 4) or that mapped discordantly (paired-reads mapping to different chromosomes or to positions separated by more than 10 times the average library size). These reads were then aligned (using Bowtie2 in --local mode to allow for soft clip alignments) to a joint TE library assembled from TE annotations^23^ belonging to 467 TE families longer than 1Kb and spanning the full range of Class I (Gypsy, Copia and LINE) and Class II TEs (MuDR, hAT and CACTA). Next, we selected all reads mapping to a TE sequence either partially (≥20nt) or fully but with an unmapped mate. These reads were re-mapped to the tomato reference genome sequence (using Bowtie2 in --local mode to allow for soft clip alignments). Read clusters composed of at least two reads mapping in the right orientation (i.e. at least one discordant read in the “+” orientation upstream of discordant read in the “-” orientation or one 3’ soft-clipped read upstream of a 5’ softclipped read, or any combination of the cases described above) were taken to indicate the presence of a bona fide nonreference TE insertion. These sites were intersected across all accessions to identify those shared and supported in at least one individual by a minimum of three reads, including at least one upstream and one downstream. Negative coverage, as defined by the minimum WGS read depth over the upstream and downstream boundaries of a putative TE insertion site, was then calculated for each accession across all putative TE insertion sites. Accessions with more than five reads negative coverage and lacking discordant- or split-reads supporting the non-reference insertion were considered as non-carriers. Accessions with less than five reads negative coverage and lacking discordant- or split-reads supporting the non-reference insertion were considered as missing information or NA.

### Validation of TIP

Visual inspection of 600 randomly chosen TIPs spanning the six TE superfamilies confirmed 82% calls, indicating high specificity of our SPLITREADER pipeline. To further assess the specificity of our bioinformatic approach we evaluated the presence of TIPs detected in the M82 cultivar on the high-quality assembled genome sequence recently obtained using Nanopore long-reads available for this accession^59^. Specifically, 1 kb sequence upstream and downstream of TIPs detected in M82 by our SPLITREADER pipeline were extracted from the Heinz 1706 reference genome (version SL2.5) and aligned using BLAT to the high-quality genome of M82. Consistent with the validation based on visual inspection, more than 70% of TIPs detected in M82 were also found in the reference M82 genome, with *COPIA* insertions showing the highest specificity (77%) (Supplementary Figure 8). Furthermore, this estimated rate is similar to the one we obtained using the same pipeline to analyze numerous *A. thaliana* resequenced genome data and which we could validate experimentally using TE sequence capture (Ref. ^27^ and Baduel et al, Methods in Mol Biol, *in press*). Using this last dataset we estimated that the false negative rate of our SPLITREADER approach is about 20%, being the sensitivity for TIPs belonging to the *COPIA*, *MuDR* and *CACTA* families the highest (Baduel et al, Methods in Mol Biol, *in press*). These rates of FP and FN are similar to those reported by others using multiple softwares developed to detect TIPs based on illumina short-reads^60^.

### SNP calling and phylogenetic analyses

Illumina re-sequencing data was aligned to the tomato genome reference v.2.50 using Bowtie2 with default parameters. The resulting alignment files were filtered to remove reads mapping to multiple locations using samtools with parameter 2-q 5 and to remove duplicated reads with Picard MarkDuplicates with default parameters (http://broadinstitute.github.io/picard, parameter REMOVE_ DUPLICATES = true). Finally, indels were realigned using GATK RealignerTargetCreator and IndelRealigner successively with default parameters Alignment files were used to call SNPs. For this, we ran GATK’s UnifiedGenotyper with default parameters in all 602 accessions simultaneously.

We extracted SNPs at 8,760 positions genotyped in the SolCAP Infinium Chip SNP microarray as indicated in the tomato annotation (ITAG2.4_solCAP.gff3). We obtained a final matrix of 1,812 SNPs after removing ambiguous SNPs and SNPs in high linkage disequilibrium using PLINK with parameters—mind 0.1—geno 0.1—indep 50 5 3.5. A phylogenetic tree was estimated from the final matrix using the ape package in R and the neighbour-joining method including *S. pennellii* LA0716 as a root. The resulting tree was plotted using the ggtree package in R. Tomato accessions in the tree were classified manually taking into account previously described classifications and their positions in the tree relative to known classifications of species and type.

### TIPs-based population differentiation

A Principal Component Analysis (PCA) using 6,906 TIPs was performed using the prcomp function in R ‘stats’ package v.3.2.3. The first two eigenvectors were retained to create a two dimensional plot.

### Genomic localization of TIPs and genes

A circos plot was constructed to represent the chromosomal distributions of genes and TEs as well as the mappability of illumina short-reads. Number of genes and TEs annotated in the reference genome as well as TIPs for the six superfamilies (Gypsy, Copia, LINE, MuDR, hAT and CACTA) were calculated in 500Kb windows using bedtools. To determine mappability we aligned Heinz 1706 short-read resequencing data (SRA: SRR1572628) on the reference genome (version SL2.5). Mappability was defined as the fraction of uniquely mapped reads (MAPQ>=10) in 10Kb windows. Gene ontology (GO) analyses were performed using AGRIGO (http://bioinfo.cau.edu.cn/agriGO/) and as input the Solyc ID of genes that contain a TE insertion within the limits of their annotation. The random expectation based on mappability bias was obtained by sampling a random set of 6906 uniquely mapped reads (MAPQ>=10) and using as input for GO analysis the Solyc ID of genes that contain uniquely mapped reads within the limits of their annotation.

### Impact of TIPs on gene expression

Raw RNA-Seq data of tomato fruit pericarp on orange stage were obtained from ref. ^35^. Expression level per gene was calculated by mapping reads using STAR v2.5.3a63 on the tomato reference genome (version SL2.5) with the following arguments – outFilterMultimapNmax 50 --outFilterMatchNmin 30 --alignSJoverhangMin 3 --alignIntronMax 50000. Duplicated pairs were removed using picard MarkDuplicates. Counts over annotations (version ITAG2.4) were normalized using DESeq2^61^. To determine the transcriptomic impact of TIPs on nearby genes (located within 1 kb), the normalized transcript levels were compared between carriers and noncarriers accessions. Analysis was restricted to 1477 genes showing expression greater than 0 in at least one sample. Variation in full-length transcripts was calculated by comparing the ratio between the normalized number of reads that mapped downstream and upstream of a given TIP. This ratio was then compared between carriers and non-carriers accessions and binned by log2 fold changes (0.5-1.5], (1.5-2.5], [3.5-4.5],>4.5.

### Linkage disequilibrium analyses

For each TIP, we calculated the pairwise r^2^ between the TIP and 300 SNPs located upstream and downstream as well as the pairwise r^2^ between all the 600 SNPs-SNPs polymorphic sites around the TIP using PLINK v2. We then contrasted the percentage of TIP-SNPs and SNP-SNPs comparisons that are in high LD (r^2^>0.4). Similar results were obtained when using r^2^>0.2 as a threshold to define polymorphisms with high LD (data not shown).

### Genome Wide Association Analysis

We recovered phenotypic information for 17 important agronomic traits in tomato, including determinate or indeterminate growth, simple and compound inflorescences, leaf morphology, fruit color, shape and taste for more than 150 accessions by web data extraction. Briefly, we performed a systematic web data scraping using google search engine (googler; https://github.com/jarun/googler) followed by text pattern matching. We noted that the World Tomato Society webpage (https://worldtomatosociety.com/) compile, in a consistent manner, phenotypic information for a large number of varieties commercialized by numerous seed-banks. We thus focused our web data scraping on this webpage. Phenotypes were either transformed in boolean (1 or 0) or quantitative variables (Supplementary Tables 2-17). For fruit color phenotype, accessions with red, purple-black or pink fruits were considered as high lycopene-containing and those with green, white, yellow or orange fruits were considered as low lycopene-containing fruits and codified as 1 or 0, respectively. Metabolomic and volatile quantitative data from ripe fruits of 397 accessions were obtained from ref. ^25,35^. SNPs and TIPs with MAF < 1% were excluded. SNP-GWAS was restricted to biallelic SNPs and LD-prunned using PLINK^62^ option --indep-pairwise 50 5 0.2. A total of 424,066 SNPs and 4,817 TIPs were retained. GWAS were performed using linear mixed models (LMM) encoded in the software EMMAX^63^. SNP-based Kinship matrix was calculated (emmax-kin-intel64 -v -d 10) and included in the models as a random effect to control population structure and minimize false positives. Manhattan and qq plots for genome wide association studies were performed using qqman R package^64^. *r*^2^ between the leading associated variant and all other associated variants in Fig. 3c were calculated using PLINK^62^ and represented by color code. Given the lower sensitivity and specificity of TIPs calling compared to SNPs, which could affect GWAS results, we investigated the robustness of our TIP-GWAS approach by randomizing 1000 times the set of carriers and non-carriers accessions and running GWAS for the two traits with validated associations (i.e. fruit color and potato-leaf)^28,42^. In both cases, the probability of detecting false associations was below the significance threshold (*P*<0.001 and *P*<0.038, respectively; Supplementary Figure 9). Association hotspots in Fig. 4a were obtained by contrasting the number of detected associations per loci with the random expectation obtained by permuting ten thousand times the localization of the detected loci across the genome and counting the number of overlapping loci. Effect sizes of associations correspond to the beta values of the leading variant from each associated loci identified by the LMM.

### Local assembly of *r*^del^ allele

Visual inspection of RNA-seq coverage of accessions harboring the *r*^del^ allele suggested a complex rearrangement. Reads from WGS of accessions carrying the *r*^del^ locus and mapping concordantly or discordantly over *PSY* locus were extracted and locally assembled using SPAdes V3.13.1^65^.

### Haplotype analysis

SNPs within 10 kb of the *PSY1* locus were retrieved for the 602 accessions and used as input into fastPHASE^66^ version 1.4.0. Default parameters were kept, except for the -Pzp option. For each SNP, haplotype membership with the highest likelihood was assigned.

### Plant materials and growth conditions

Tomato seeds from S. lycopersicum cv. Heinz 1706 and TS-666 were grown in a grow chamber (Percival) using 20 l pots, 16/8 h photoperiod, 24±3 °C, 60% humidity, and 300±100 mmol m^-2^ s^-1^ incident irradiance. Pericarp for at least four ripe fruits from two plants were harvested 60 days after anthesis, immediately frozen in liquid N2 and kept at −80 °C until use.

### Full-length cDNA nanopore sequencing

Total RNA was extracted from 100 mg of ripe fruits using the Nucleo-spin RNA Plant mini kit (Macherey-Nagel). Library preparation and Nanopore sequencing were performed at the Ecole normale superieure genomic core facility (Paris, France). After checking RNA quality by Fragment Analyzer, 10 ng of total RNA were amplified and converted to cDNA using SMART-Seq v4 Ultra Low Input RNA kit (Clontech). 17 fmol of cDNA was used for library preparation using the PCR Barcoding kit (SQK-PBK004 kit; ONT) and cleaned-up with 0.6X Agencourt Ampure XP beads. 2 fmol of the purified product was amplified during 18 cycles, with a 17 minutes elongation step, to introduce barcodes. Samples were multiplexed in equimolar quantities to obtain 20 fmol of cDNA and the rapid adapter ligation step was performed. Multiplexed library was loaded on an R9.4.1 flowcell (ONT) according to the manufacturer’s instructions. An standard 72-hour sequencing was performed on a MinION MkIB instrument. MinKNOW software (versions 19.12.5) was used for sequence calling. Long-reads were mapped on the tomato reference genome (SL2.5) using minimap2^67^ V2.11-r797 and visualized with Integrative Genomic Viewer.

## DATA AVAILABILITY

Long-read nanopore sequencing data has been deposited in the European Nucleotide Archive (ENA) under project PRJEB37834 [https://www.ebi.ac.uk/ena/data/view/PRJEB37834]. A reporting summary for this Article is available as a Supplementary Information file. The datasets generated and analyzed during the current study are available from the corresponding author upon request. The source data underlying Fig. 1a, 1c, 1d, 2c, 2d, 3f, 3g, 4a, 4b, 4d, 5c, 5e and 5g are provided as a Source Data file.

## CADE AVAILABILITY

Codes used to detect TE insertions are available at https://github.com/LeanQ/SPLITREADER with no restriction to access.

## Acknowledgments

We thank members of the Colot group and especially Pierre Baduel for discussions and critical reading of the manuscript. We thank Zachary Lippman for sharing his high-quality genome assembly of M82 before publication. We also thank the World Tomato Society for making available phenotypic information of tomato cultivars. Support was from the Agence National de la Recherche (ANR-17-tomaTE to V.C. and J.J.G.), the Centre National de la Recherche Scientifique (MOMENTUM program, to L.Q.) and France Génomique national infrastructure, funded as part of the “Investissements d’Avenir” program managed by the Agence Nationale de la Recherche (contract ANR-10-INBS-09)

## Author contributions

J.J.-G., V.C and L.Q conceived the project. E.D. and L.Q. performed the detection of TIPs. J.J.-G. performed SNP calling. M.B, B.L. and L.Q. performed ONT experiments. M.D. and L.Q. analyzed the data. V.C. and L.Q. wrote the manuscript with additional input from M.D. All the authors read and approved the manuscript.

## SUPPLEMENTARY FIGURES

**Supplementary Figure 1.**
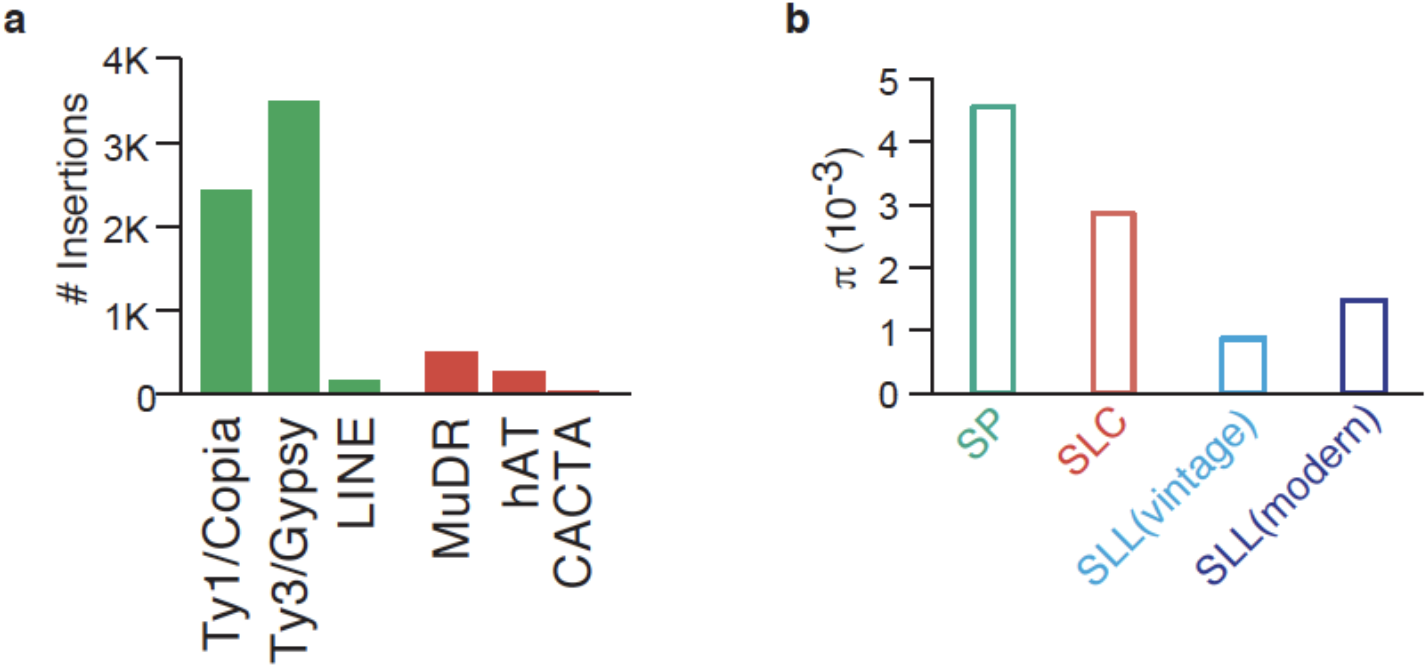
The tomato mobilome and genetic diversity per group. **a.** Number of TIPs per TE superfamily, in green Class I LTR and non-LTR retroelements, and in red DNA transposons. **b**.Genetic diversity per tomato group.

**Supplementary Figure 2.**
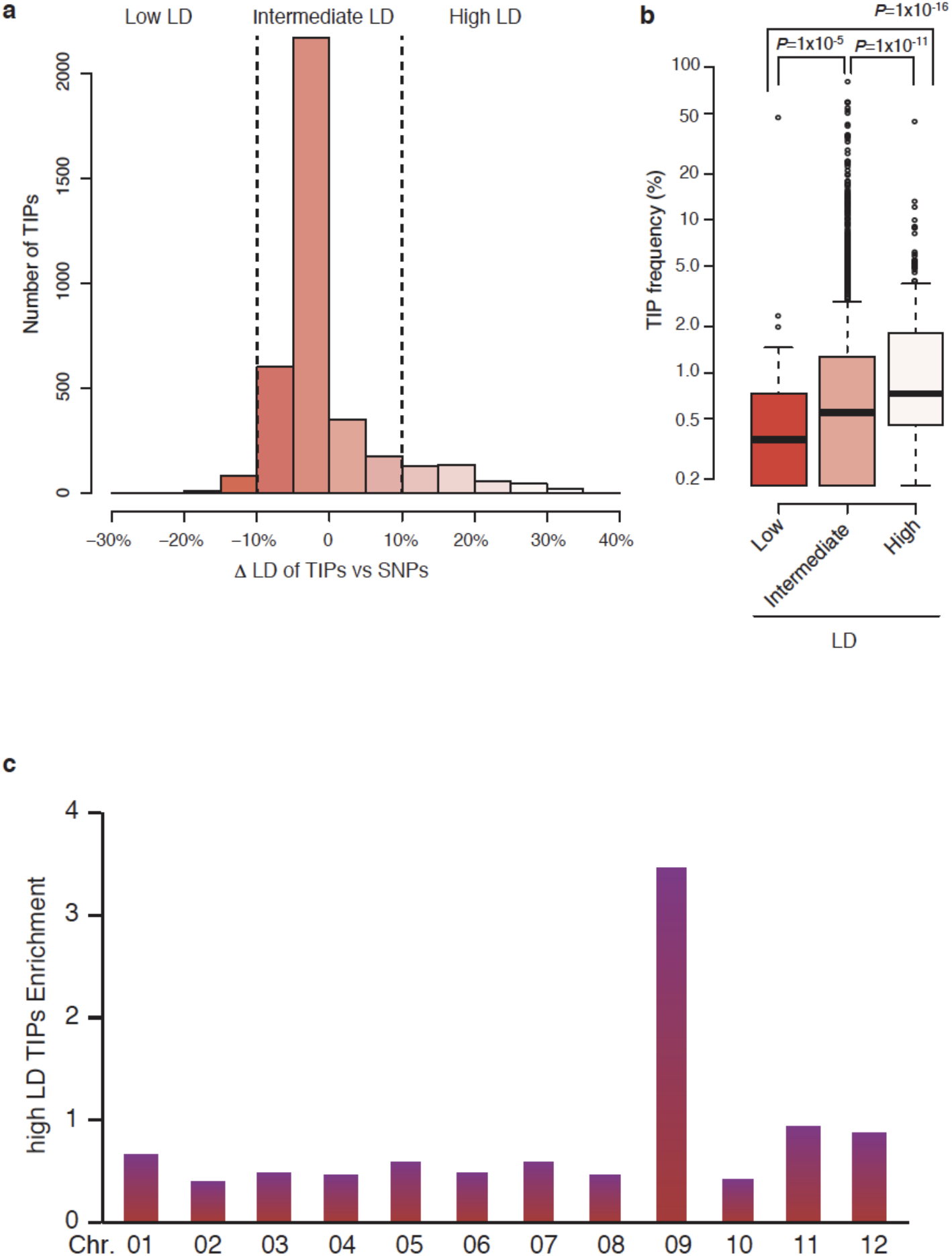
LD between TIPs and SNPs. **a.** Distribution of the proportion of SNPs that are in lower or higher linkage disequilibrium (LD) with TIPs or other SNPs. **b**. LD between TIPs and SNPs in relation to TIPs frequency. For each boxplot, the lower and upper bounds of the box indicate the first and third quartiles, respectively, and the center line indicates the median. Statistical significance for differences was obtained using the MWU test. **c**. Enrichment of high LD TIPs per chromosome

**Supplementary Figure 3.**
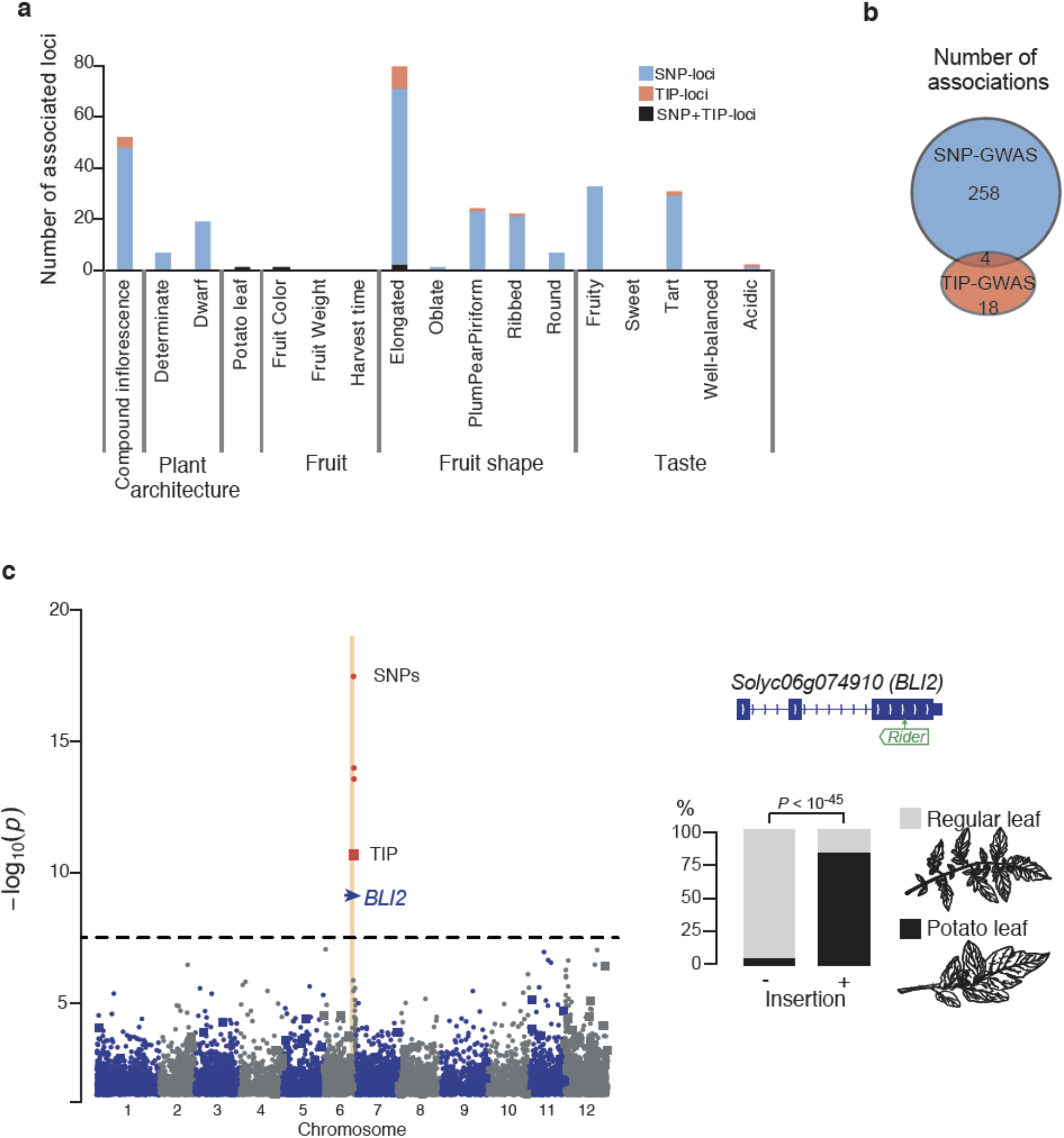
TIP- and SNP-GWAS of agronomically important traits. **a.** Number of loci associated with variation in 17 traits. loci detected by both SNPs and TIPs are indicated in black. **b.** Significant associations detected by SNP- and TIP-GWAS and their overlap. **c**. SNP- and TIP-based GWAS (circles and squares, respectively) results for potato-leaf.

**Supplementary Figure 4.**
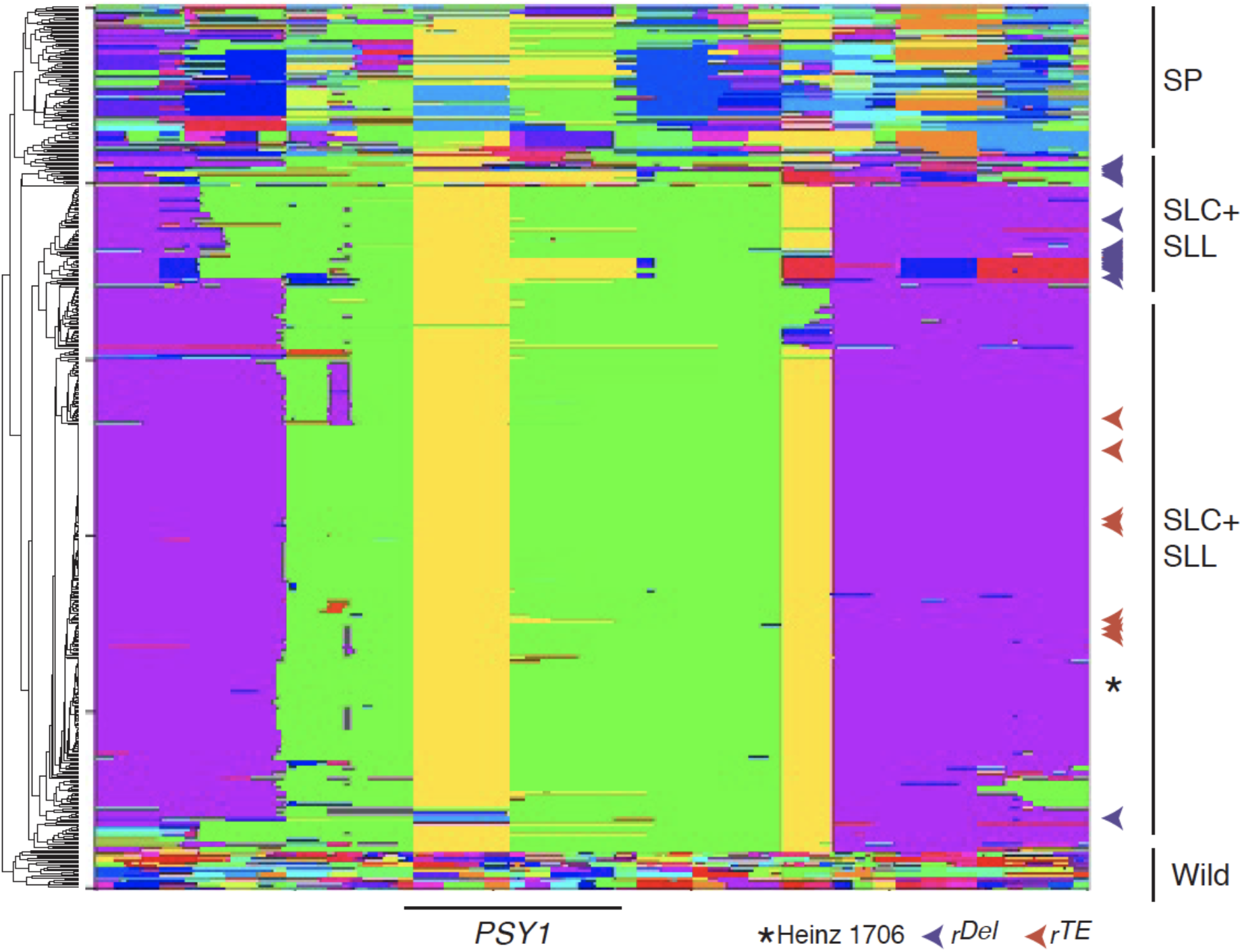
Reconstruction of *PSY1* haplotypes.

**Supplementary Figure 5.**
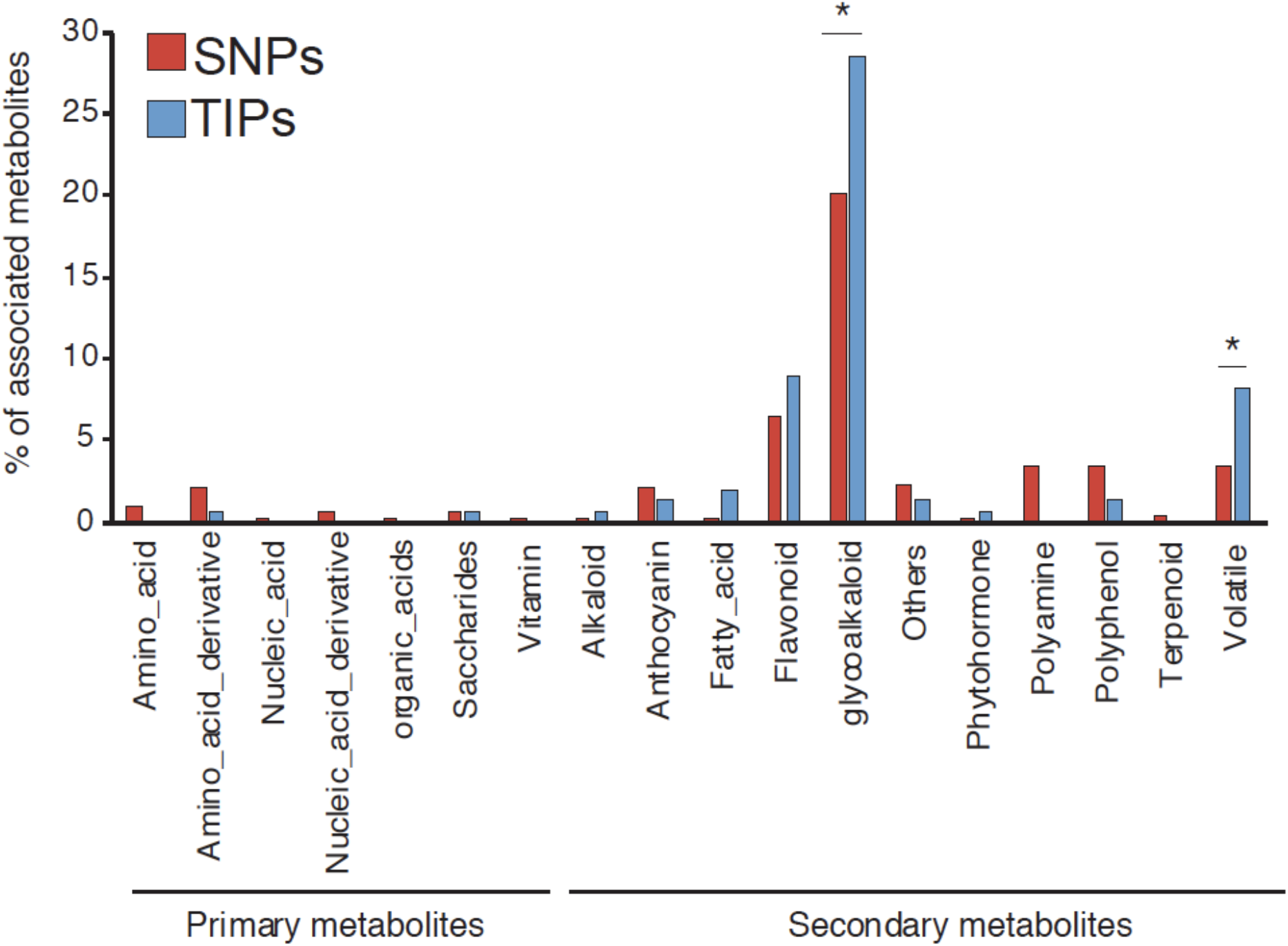
Percentage of loci associated with distinct metabolites. Statistical significance for differences was obtained using the Fisher test. * indicates p-value <0.05

**Supplementary Figure 6.**
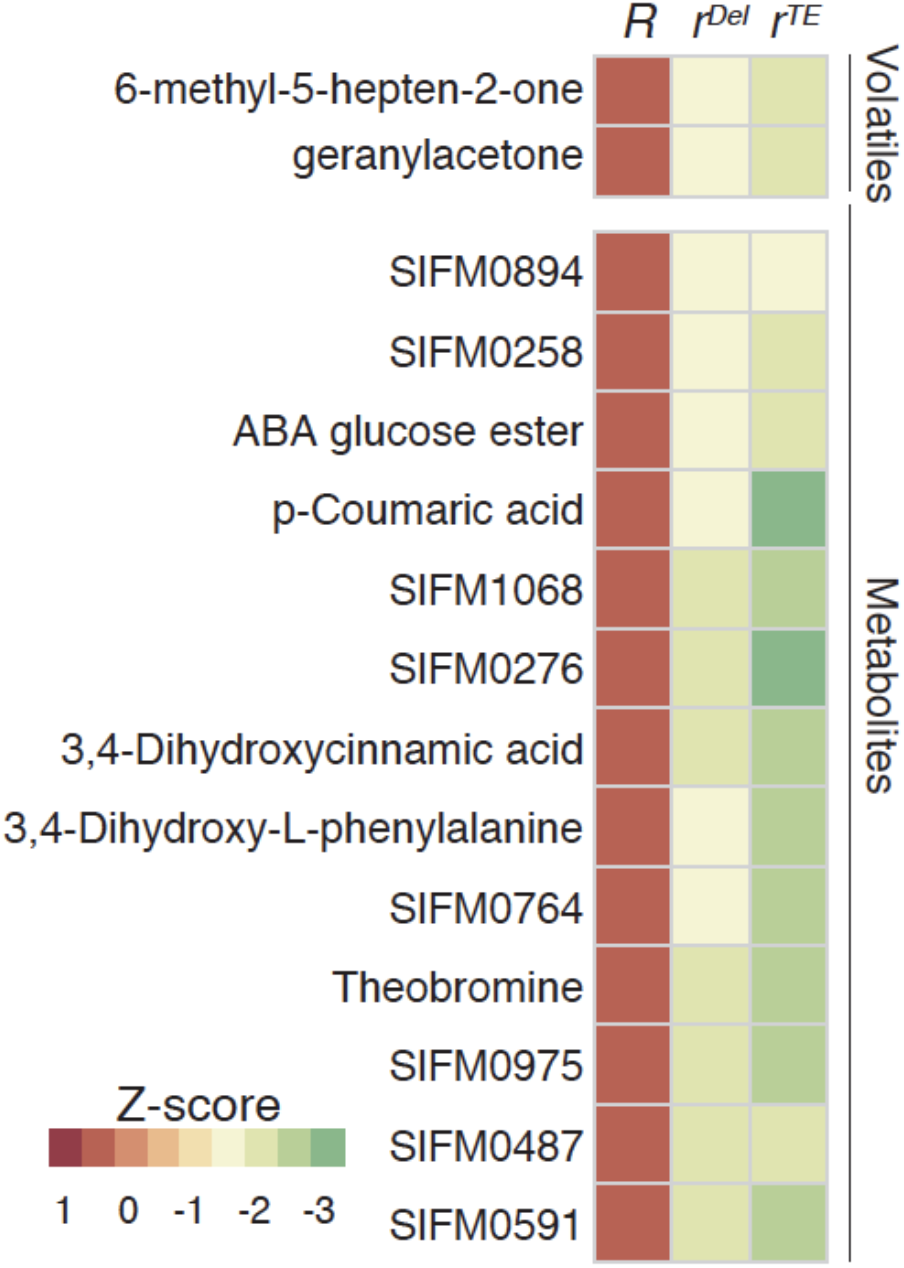
Normalized levels of 15 metabolites associated with *r^del^* and *r^TE^*.

**Supplementary Figure 7.**
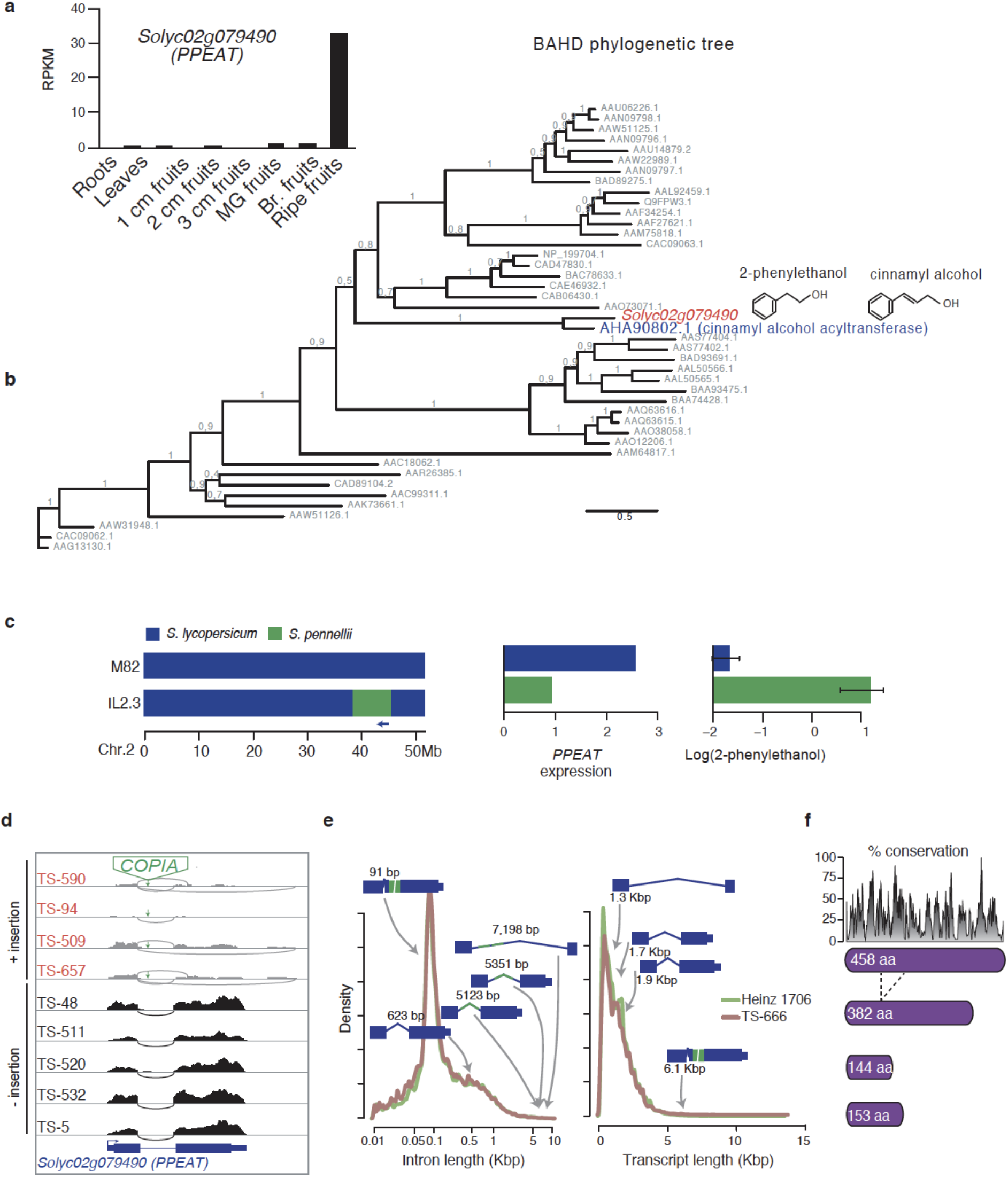
*Solyc02g079490* encodes a putative 2-phenylethanol Acyl-CoA transferase. **a**. Expression profile of *Solyc02g079490* in leaves and during fruit development. **b.** phylogenetic tree of the characterized plant BAHD family of acyltransferases. **c**. Genotype, *Solyc02g079490* expression and 2-phenylethanol levels of selected ILs and M82 control. **d**. Genome browser view of RNA-seq coverage over *Solyc02g079490* of accessions carrying or not the associated TE insertion. **e**. Intro and transcript length distribution based on our Nanopore sequencing. *PPEAT* transcripts are indicated. **f**. % conservation of PPEAT based on the alignment of 45 plant BAHD acyltransferases (from b).

**Supplementary Figure 8.**
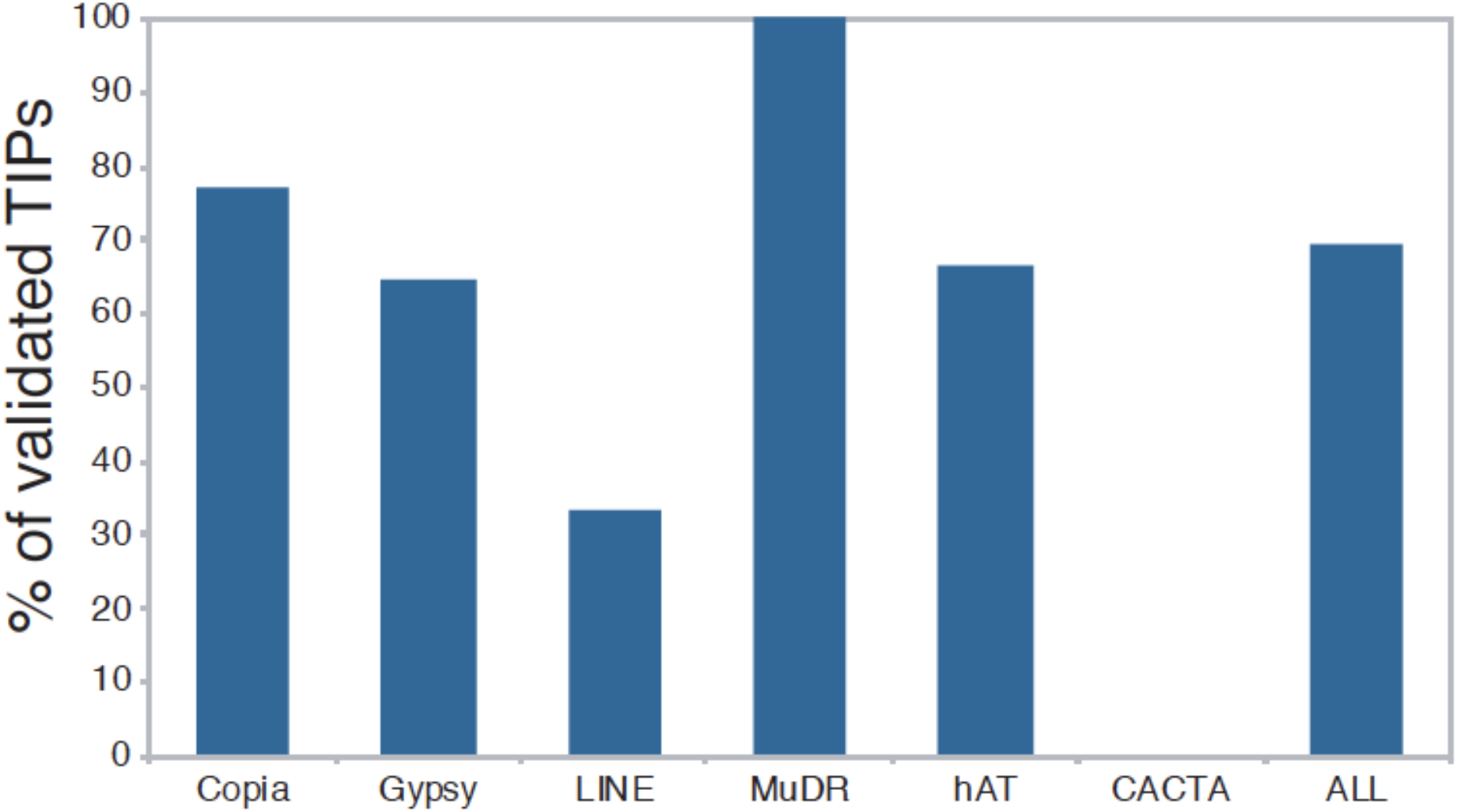
Specificity of the SPLITREADER pipeline.

**Supplementary Figure 9.**
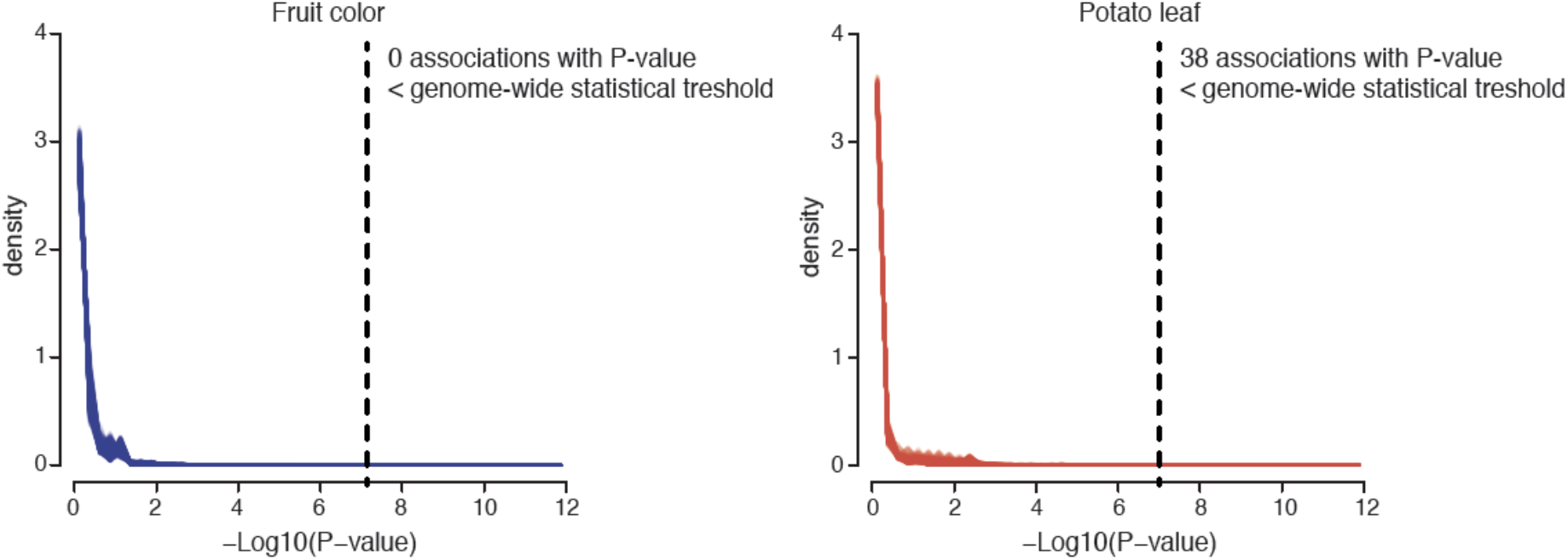
Empirical distribution of expected TIP-GWAS P-values under the null hypothesis generated by randomizing genotypes:traits pairs 1000 times.

